# Molecular dynamics simulations of membrane deformation induced by the amphiphilic helices of Epsin, Sar1p and Arf1

**DOI:** 10.1101/140970

**Authors:** Zhen-lu Li

## Abstract

The N-terminal amphiphilic helices of proteins Epsin, Sar1p and Arf1 play a critical role in initiating membrane deformation. We present here the study of the interactions of these amphiphilic helices with the lipid membranes by combining the all-atom and coarse-grained simulations. In the all-atom simulations, we find that the amphiphilic helices of Epsin and Sar1p have a shallower insertion depth into the membrane compared to the amphiphilic helix of Arf1, but remarkably, the amphiphilic helices of Epsin and Sar1p induce higher asymmetry in the lipid packing between the two monolayers of the membrane. The insertion depth of amphiphilic helix into the membrane is determined not only by the overall hydrophobicity but also by the specific distribution of polar and non-polar residues along the helix. To directly compare their ability of deforming the membrane, we further apply coarse-grained simulations to investigate the membranes deformation under the insertion of multiple helices. Importantly, it is found that the amphiphilic helices of Epsin and Sar1p generate a larger membrane curvature than that of Arf1, in accord with the experimental results qualitatively. These findings enhance our understanding of the molecular mechanism of the protein-driven membrane remodeling.

## I INTRODUCTION

The transport of cargo molecules in both endocytic and secretory pathways is conducted by coated vesicles. The formation of these vesicles involves many membrane remodeling proteins, such as the well-known coat proteins including clathrin in endocytic vesicle and contator in COPII and COPI transport vesicle.^1–3^ However, clathrin, COPII or COPI are not directly in contact with the membrane. Instead, there is a kind of proteins which anchor to the membrane directly and can trigger an initial membrane deformation. Then, they further recruit membrane remodeling proteins (such as clathrin, COPII or COPI) from the cytoplasm to generate membrane invagination or budding. Epsin is a representative one of such proteins, and is involved in the clathrin-mediated endocytic pathways.^4^ Similarly, upon exchange of GDP to GTP, small GTPase Sar1p and Arf1 initiate the membrane bending during COPII and COPI transport pathway separately.^5–7^

The localization of Epsin, Sar1p or Arf1 at the membrane is achieved by embedding its short N-terminal amphiphilic helix shallowly into one leaflet of the lipid membrane. *In vitro* experiments revealed that the association of Epsin, Sar1p or Arf1 with the membrane could deform the membrane. For example, Epsin could convert PI(4,5)P_2_ containing-liposomes into tubules with a diameter of 19 nm.^8^ In the GTP-bound state, Sar1p could convert liposomes into tubules with a mean diameter of 26 nm,^9^ and the myristoylated Arf1 could transform negatively-charged liposomes into tubules of 45 nm in diameter.^10^ All these proteins show the ability to curve membrane into a bud or a tube with the size of a few tens of nanometers both *in vitro* and *in vivo* experiments. Thus it is very important to clarify how these proteins deform the membrane, which could help better understand the generation of cellular traffic vesicles.

It is known that the N-terminal amphiphilic helices of proteins like Espin, Sar1p and Arf1 play a crucial role in inducing membrane deformation.^11–14^ The N-terminal amphiphilic helix is embedded into the lipid matrix, with their hydrophobic face inserting between fatty acyl-chain and their polar face exposing to the water. The so-called local spontaneous curvature mechanism (or hydrophobic inserting mechanism) reveals that a shallow insertion of an amphiphilic helix into only one leaflet of the membrane can perturb the packing of the lipid polar headgroups and induce a local monolayer deformation.^11,12^ Theoretical work by Campelo *et al*. showed that the insertion of amphiphilic helices can be very powerful in bending the membrane, and they also indicated that shallow insertions are best suited for the production of high membrane curvature.^15^

In the computational study of peptide-membrane interaction, molecular dynamics simulation is a very useful tool as it provides detailed structural and dynamical information.^16–20^ So far, many simulation works have focused on the N-BAR domain, as the dimer of this protein has both a charged, crescent-shaped surface and two amphiphilic helices, which may bend the membrane through combining a scaffold mechanism and the hydrophobic inserting mechanism.^21–26^ Recently, the membrane remodeling effects of anti-bacteria peptides^27^ and monomeric *α*-Synuclein^28,29^ that are involved in the mitochondrial remodeling were also studied. However, to the best of our knowledge, the role of these amphiphilic helices of Epsin (M_1_STSSLRRQMKNIVHN_16_), Sar1p (M_1_AGWDIFGWFRDVLASLGLWNKH_23_) and Arf1 (G_2_NIFANLFKGLFGKK_16_) in generating membrane deformation have not been investigated in previous computational studies. Actually, though participating in rather different cellular transport processes, Epsin, Sar1p and Arf1 share a similar mechanism that upon stimulus of signaling lipids or nucleotide, the N-terminal segment of Epsin, Sar1p or Arf1 folds into an amphiphilic helix and inserts into the membrane. Therefore, it is also of great importance to reveal the similarity and difference between their interactions with cell membrane.

In the present work, we will study the molecular mechanism of the amphiphilic helices of Epsin, Sar1p and Arf1 deforming the membrane by combining the all-atom and coarse-grain simulations. In particular, we will investigate how the membrane bending ability of different amphiphilic helices are related to their detailed physicochemical properties. The all-atom simulations are performed to investigate the complex of a single helix associated with the membrane. Particularly, the influence of the helix insertion on the lipid packing of the membrane are analyzed. The coarse-grained simulations at a large scale are performed in order to directly illustrate the membrane deformation process under the insertion of multiple helices into the membrane.

## II MODEL AND METHODS

In the all-atom (AA) simulations, a membrane of 84 DOPC (1,2-Dioleoyl-sn-glycero-3-phosphocholine) and 36 DOPS (1,2-di-(9Z-octadecenoyl)-sn-glycero-3-phospho-L-serine) molecules is prepared (certain amounts of anionic lipids (30%) are added into the membrane, as did in several previous studie^24–28^). The membrane is equilibrated for 100 ns before further application. The N-terminal helices of Epsin, Sar1p and Arf1 are equilibrated for 10 ns in solvents respectively, and the final configurations are shown in Fig. 1a. The corresponding helical wheel representations for these helices are shown in Fig. 1b. Next, the helix is pre-inserted into the membrane by using the InflatGro method,^30,31^ with its hydrophobic face interacting with the membrane interior while the hydrophilic face interacting with the water. In order to enhance simulation sampling, for each kind of helix of Epsin, Sar1p and Arf1, two different initial simulation configurations of the helix relative to the membrane are applied, by displacing the helix with its long axis either parallel to the x-axis or parallel to the y-axis of the simulation box. Water molecules are then added to the system. The charges of DOPS molecules and the amphiphilic helix are neutralized by adding appropriate number of counter-ions. A restrained NVT equilibrium simulation (the mainchain C_*α*_ atoms are restrained) of 1 ns and a followed restrained NPT simulation of 40 ns are performed to relax the simulation system. The unstrained run is performed for 360 ns for analysis.

**Figure 1.**
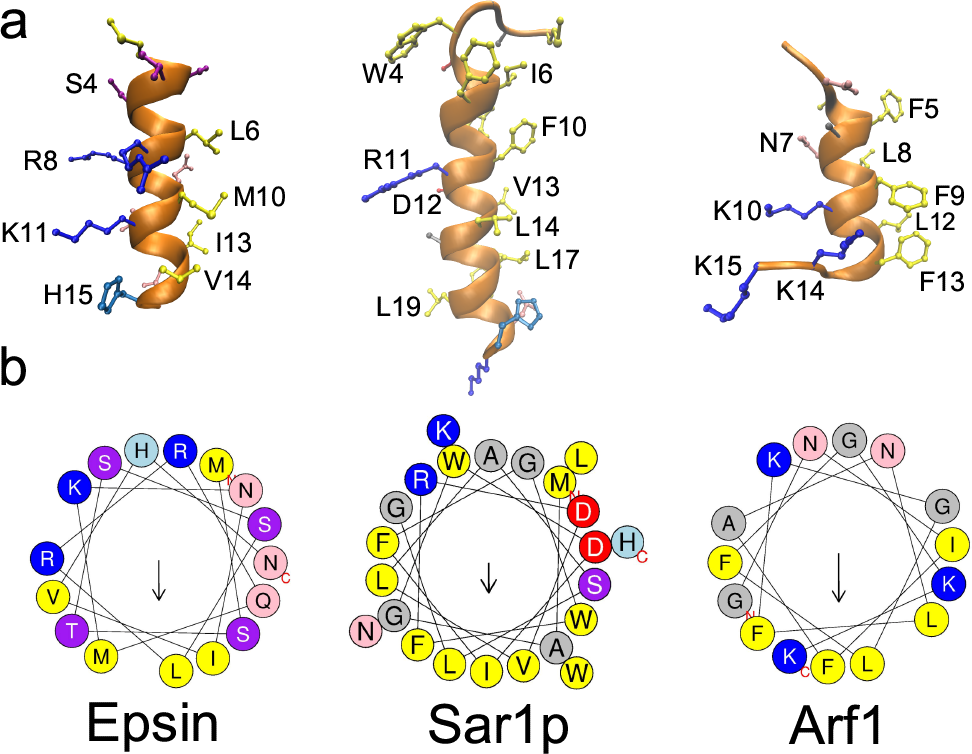
(a) The configurations and (b) the corresponding helical wheel representations of the amphiphilic helices of Epsin, Sar1p and Arf1. Color coding for residues: yellow, hydrophobic; purple, serine and threonine; blue, basic; red, acidic; pink, asparagine and glutamine; grey, alanine and glycine; green, proline; light blue, histidine. The arrow in helical wheels corresponds to the hydrophobic moment.

The CHARMM36 all-atom force field is used for the phospholipids as well as the amphiphilic helix (the histidine is in neutral form).^32,33^ The TIP3P model is used for water.^34^ The electrostatic interactions are treated by Particle-Mesh Ewald (PME) method^35^ with a real-space cutoff of 1.2 nm. The van der Waals interactions are also cut at 1.2 nm. Bond lengths are constrained via the P-LINCS algorithm.^36^ A time step of 2 fs is employed. The center of mass motion is removed in every step, and the neighbor list is updated in every 10 steps. The temperature is coupled by using Nose-Hoover thermostat at 310 K,^37,38^, whereas the pressure control is achieved by semi-isotropic Parrinello-Rahman scheme at 1 bar.^39^

The MARTINI coarse-grained force field (including the improved protein force filed) which is developed by Marrink’s group is applied for the coarse-grained (CG) simulations.^40–42^ In the improved MARTINI protein force field, the polarized water is used and the residues are reparametrized through reassignment of bead types or by introducing embedded charges. These procedures make the force field better to describe the peptide-membrane interactions. The Lennard-Jones (LJ) potentials are cut off at a distance of 1.2 nm and smoothly shifted to zero between 0.9 and 1.2 nm. The electrostatic interactions are also cut off at a distance of 1.2 nm with smooth switching of the interactions from 0.0 to 1.2 nm. The relative dielectric constant is set as 1. A large rectangular membrane consisted of 3024 DOPC and 1296 DOPS is prepared, and the simulation box is about 110 nm in X-axis direction and about 14 nm in Y-axis direction. Twenty helical peptides are pre-inserted into the membrane with their long axis along with the Y direction.^24^ The interval space between helices is about 2 nm along the X direction. About 190000 polarized water molecules are added to the system, and counter-ions are included to neutralize the charge of DOPS and helices separately. The coarse-grained simulations are performed in the NPT ensembles coupling to a bath with a constant temperature (*T*) of 310 K and a constant pressure (*P*) of 1 bar.^43^ A time step of 20 fs is used in the CG simulations. Notice that the effective time sampled in CG simulations is 4 times as large as that in atomistic simulations,^44^ so here the effective simulation time step is approximately 80 fs. For each CG simulation, a simulation of 2 *µ*s is performed.

All the all-atom and coarse-grained simulations and the analysis are performed by using GROMACS 4.6.5 software package.^45^

## III RESULTS AND DISCUSSION

### A. Partitioning of amphiphilic helices in the membrane

Fig. 2 shows the final configurations of the N-terminal helices of Epsin, Sar1p and Arf1 binding to the membrane as well as the density distribution of the lipid (P or C2 atoms) and the helical peptide (backbone atoms). The helices are roughly parallel to the membrane surface and embedded into the lipid matrix partially. The center of mass of the backbone atoms of these helices is at the level of the lipid glycerol group. Specifically, for Epsin and Sar1p, the amphiphilic helix is distributed between the lipid glycerol group (C2) and phosphate group (P), while for Arf1, it is below the average position of glycerol group (C2). Fig. 3a further shows the time evolutions of the distance of these amphiphilic helices to the membrane center. Starting from a similar initial displacement, these helices gradually adjust themselves to their most preferred distribution. As shown in Table. I, the averaged insertion depth of the helix of Arf1 is the deepest (about 1.52 nm from the membrane center), while the N-terminal helices of the Epsin and Sar1p have shallower insertion depth of about 1.96 and 1.83 nm respectively.

**Figure 2.**
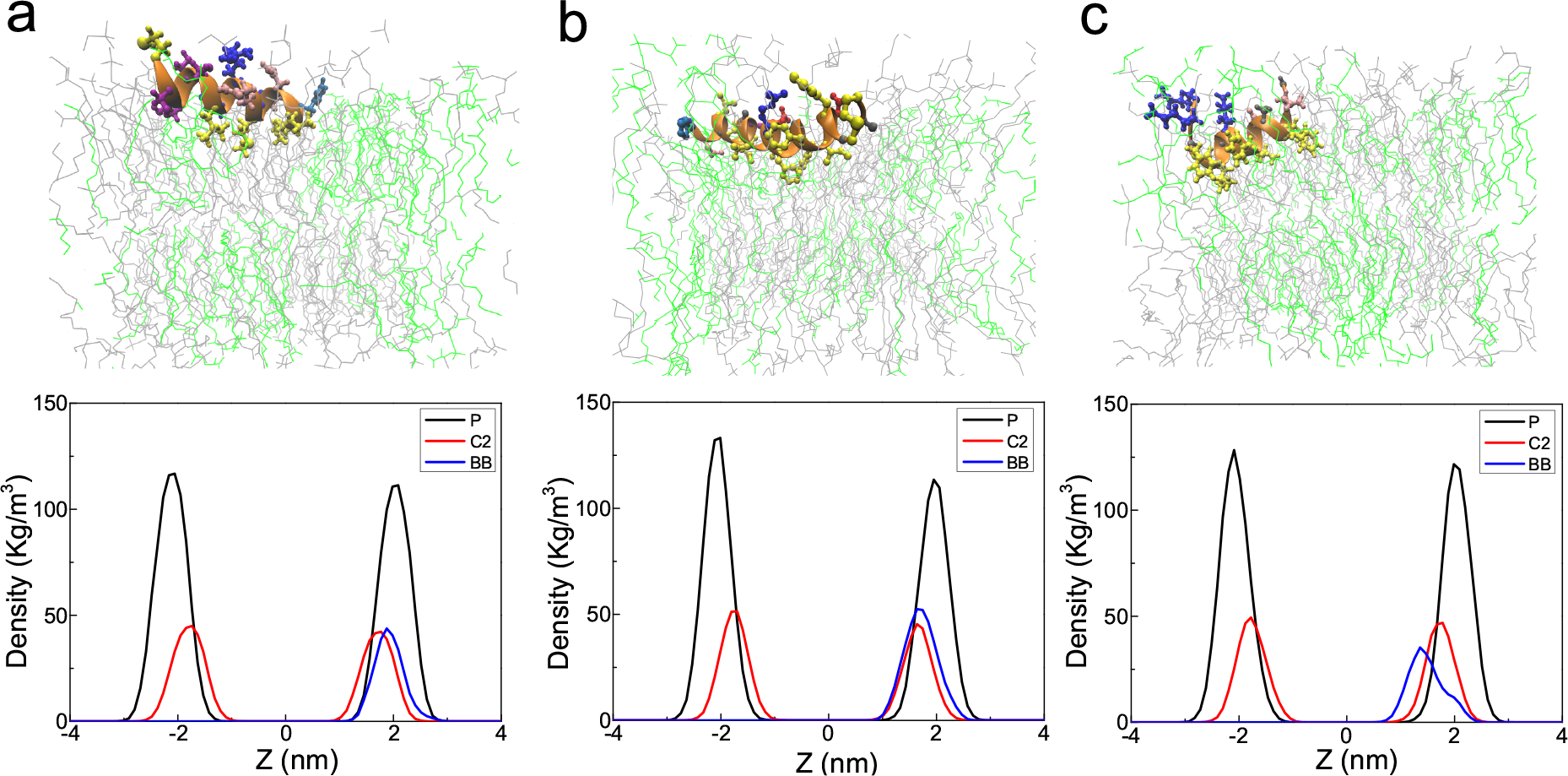
(a) Top: Final configuration of N-terminal helix of Epsin, Sar1p and Arf1 binding to the membrane. DOPC is in grey and DOPS is in green. The color code for the helix is same as Fig. 1.(b) Bottom: The density distribution of lipid P and C2 atoms, and of the backbone (BB) atoms of the helix.

**Figure 3.**
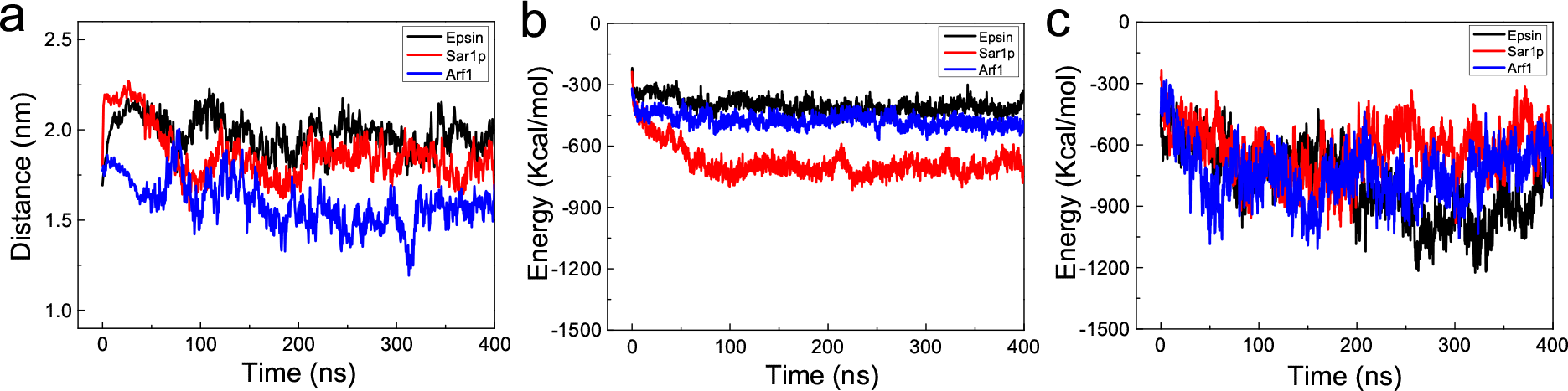
(a) Time evolutions of the distance between the center of mass of the backbone atoms of the helices of Epsin, Sar1p and Arf1 and the center of mass of lipid membrane along the Z axis.(b) and (c) Time evolutions of the Van der Waals and the electrostatic interaction between the lipid membrane and the helices of Epsin, Sar1p and Arf1.

**Table I.** Properties of the amphiphilic helices of Epsin, Sar1p and Arf1: L for sequence length, z for the net charge, H for the hydrophobicity and *µ*H for the hydrophobic moment. *D_h_* denotes the insertion depth of the helix in the all atom simulations. *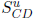* and *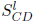* denote the order parameter of the upper and lower leaflets in the all atom simulations (averaged over the last 200 ns). R denotes the estimated curvature radius of the membrane deformation induced by the insertion of multiple helices in the CG simulations. This value is took averaged over the last 1 *µ*s simulation.

| Helix | L | z(e) | H | $\mu H$ | $D_h(nm)$ | $S_{CD}^u$ | $S_{CD}^l$ | R(nm) |
| --- | --- | --- | --- | --- | --- | --- | --- | --- |
| Epsin | 16 | +3 | 0.131 | 0.483 | $1.96 \pm 0.08$ | 0.136 | 0.162 | $59 \pm 6$ |
| Sar1p | 23 | 0 | 0.707 | 0.270 | $1.83 \pm 0.08$ | 0.125 | 0.162 | $55 \pm 8$ |
| Arf1 | 15 | +3 | 0.447 | 0.424 | $1.52 \pm 0.09$ | 0.149 | 0.160 | $152 \pm 68$ |

According to the experimental results, Epsin and Sar1p initiate membrane tubes with smaller diameter (compared to Arf1). So possibly a deep insertion depth of the amphiphilic helix into the membrane is not suited for bending the membranes. Instead, a shallow insertion depth into the membrane can be more efficient in creating high membrane curvature. This point is also indicated in several theoretical works on the short peptide interacting with the membrane.^15,48^ In particular, theoretical work by Campelo *et al*. showed that, for a membrane with monolayer thickness of 2.0 nm and 2.2 nm respectively, the optimal insertion depth of the helix (where the helix can initiate the largest membrane spontaneous curvature) is about 1.75 nm and 1.9 nm from the membrane center. Here, in our simulations, the thickness of membrane monolayer (measured from the peaks of the phosphate distribution) is about 2.05 nm. Based on the theory prediction, a rough estimate of the optimal insertion depth is around 1.8 nm for the membrane studied here. The insertion depth of the helix of Arf1 is 1.52*±*0.09 nm, much lower than 1.8 nm. The insertion depth of the N-terminal helices of Epsin and Sar1p are 1.96*±*0.08 nm and 1.83*±* 0.08 nm respectively, which are very similar with the theoretical prediction. We should notice that he peptide is usually simplified as a simple cylinder in the theoretical model, while in the molecular simulation, it considers the specific structures at residue level. Besides, in the simulation, the position of these helices also undergoes essential fluctuation. These factors may lead to discrepancies between the value of theoretical prediction and molecular simulation. Nevertheless, the results of the theoretical prediction and molecular simulations are comparable—a deep insertion of the helix into the membrane slightly contribute to the membrane deformation, while a shallow insertion of the helix at the level of (or slightly above) membrane hydrophilic-hydrophobic interface is suitable for bending the membrane. Taken together the theoretical and simulation results, we suggest that an insertion depth around 1.75 nm to 2.0 nm should be very suited for the production of high membrane curvature.

### B. Factors that influence the insertion depth of the amphiphilic helices into the membrane

Fig. 3b-c show the pair interactions between the amphiphilic helix and the membrane. The Van der Waals interaction between the helix of Sar1p and the membrane is the lowest. The value is almost 2 times and 1.5 times lower than the interactions between the membrane and the helix of Epsin or the helix of Arf1 respectively. This is partly due to the longer sequence length of the helix of Sar1p (1.4 times larger than the helix of Epsin and 1.5 times larger than the helix of Arf1; see Table. I). More importantly, the high hydrophobicity of Sar1p considerably contributes to the low Van der Waals interaction. In contrast, the electrostatic interactions of the helices of the Epsin and Arf1 with the membrane are much lower, compared to the helix of Sar1p. This is because both the helices of Epsin and Arf1 have three net positive charges, while the helix of Sar1p has no net charge. Therefore, the former two helices will have stronger electrostatic interactions with the anionic DOPS lipid molecules.

A high hydrophobicity of peptide is favorable for the strong binding to the membrane and a deep insertion depth into the membrane. However, the insertion depth also depends on the interaction of polar and charge residues with the lipid headgroup and water.^46^ Fig. 4 shows the insertion depth of individual residues into the membrane. For each amphiphilic helix, the residues that have a relatively deep insertion into the membrane are mostly hydrophobic residues. These residues include L6, M10, I13 and V14 in the helix of Epsin, residues I6, F7, W9, F10, V13, L14, L17 in the helix of Sar1p and residues F5, L8, F9, L12, F13 in the helix of Arf1. Typically, residues at the bottom of the helix wheel in Fig. 1 (the hydrophobic face) have deep insertion into the lipid acyl chains, in exception that K15 and K16—at the end of the helix of Arf1, are at lipid-water interface due to their favorable interactions with the lipid headgroup. In contrast with the hydrophobic residues, polar and charge residues prefer to stay at the lipid-water interface, driven by the polar as well as electrostatic interaction with the lipid headgroup or water. Hydrogen bonds formed between the residues and the lipids are shown in Fig. 4. Remarkably, positively charged residues including arginine and lysine are largely involved in the hydrogen bonds interaction with the lipid molecules (typically 1-2.5 hydrogen bonds for each). Polar residues also form hydrogen bonds with the lipid molecules, but at a less extent.

**Figure 4.**
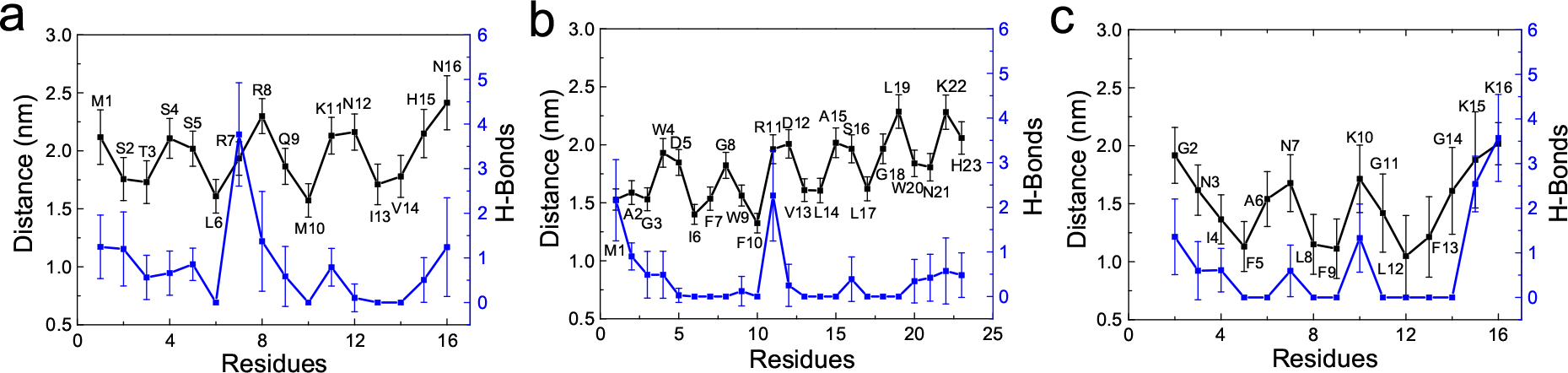
The average insertion depth of individual residues into the membranes (black line) and the number of hydrogen bonds formed between residues and the lipid membranes (blue line). (a-c) are for the N-terminal helices of Epsin, Sar1p and Arf1 respectively.

The distribution of polar and non-polar residues along the helix is of great importance in the insertion of the amphiphilic helix into the membrane. Supposing that an overall hydrophobic face has few polar residues incorporated (such as N20 in Sar1p), a deep insertion of the amphiphilic helix into the membrane will become unfavorable since the polar residues are unlikely to interact with the hydrophobic acyl chains of lipid. Regarding the amphiphilic helix of Sar1p, the hydrophobic and hydrophilic face of the helix are mixed with few polar and non-polar residues respectively. Even though the helix of Sar1p has a high hydrophobicity, the helix only has a modest insertion depth into the membrane. To quantitatively depict the the amphiphilicity of a helical peptide, we introduce the hydrophobic moment (*µ*H)^47^ that tells whether a helix exhibits clearly one hydrophobic face and one polar face. A large *µ*H value means that the helix is amphiphilic perpendicular to its axis. Though being highly hydrophobic, the helix of Sar1p only has a low *µ*H due to the doping of polar residues at its hydrophobic face and nonpolar residues at its hydrophilic face. The helix of Arf1 has medium hydrophobicity and *µ*H (see Table I). Excluding the two lysines (K15K16) at the end of the helix of Arf1, the hydrophobic face of the helix of Arf1 is wide and clear. These features contribute to the deepest insertion depth into the membrane for the helix of Arf1. The helix of Epsin has large *µ*H but very low hydrophobicity. The hydrophobic face of the helix of Epsin is very small, therefore it only has a shallow insertion depth into the membrane.

### C. Effect of amphiphilic helices on the lipid packing of the membrane

The insertion of the helix will influence the lipid packing of membrane. This effect can be studied by calculating the deuterium order parameter (S_*CD*_) of the lipid acyl chain. For each amphiphilic helix, the order parameter is shown in Fig. 5a and Fig. 5b for the upper leaflet (containing the helix) and the lower leaflet respectively. The order parameter for the upper leaflet in the present of N-terminal helices of Epsin and Sar1p decreases a lot compared with the order parameter for a pure membrane, while for the helix of Arf1, the value nearly has no change in spite of its deep insertion into the membrane. Oppositely, as for the lower leaflet without an embedded helix, the order parameter has an approximately equal increase for each system.This indicates that the insertion of the helix into one leaflet has non-negligible impact on the opposite leaflet of the membrane. This is reasonable considering the coupling between the membrane’s two leaflets—the opposite leaflet makes corresponding adjustment to the changes in the packing of acyl chains of the upper leaflet. Thus, the asymmetry in the order parameter of the two leaflets could be an important driving force for the membrane deformation.^29,49^ The asymmetry in the lipid packing of the two leaflets probably implies an asymmetry in the stress pressure for the two leaflets, and an asymmetric stress pressure profile results in generation of membrane deformation.

**Figure 5.**
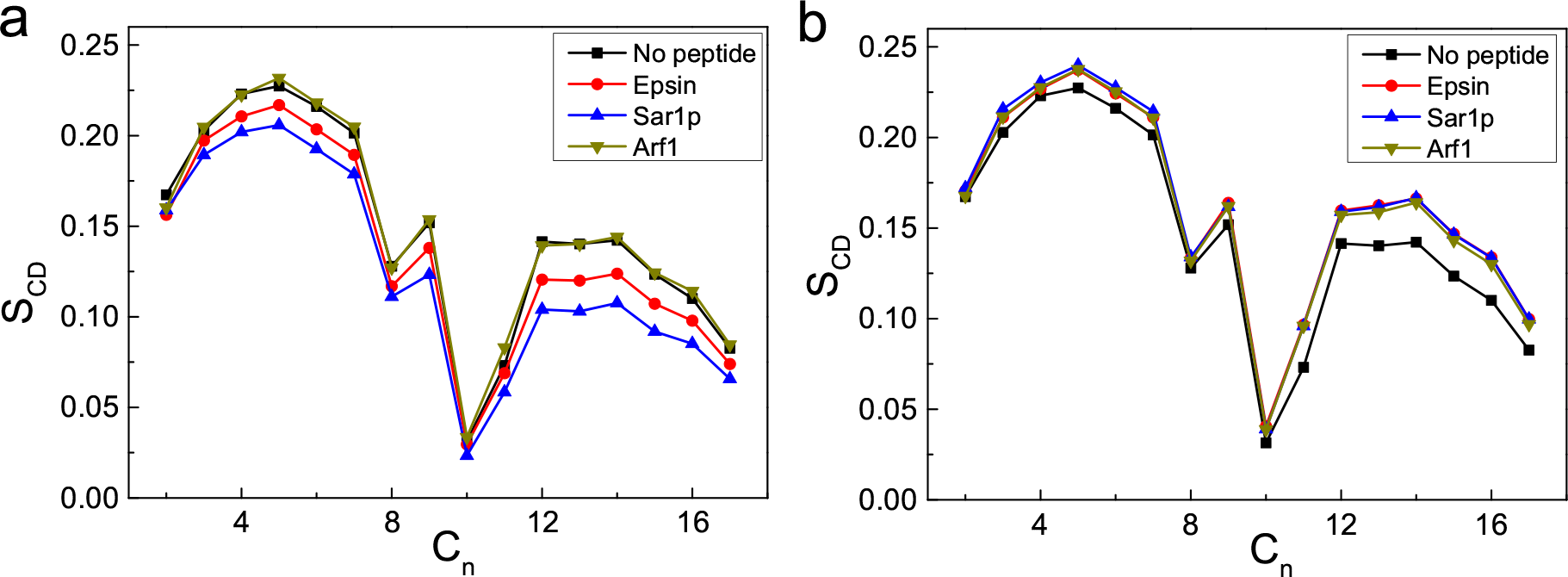
The deuterium order parameter of the lipid acyl chain for (a) the upper leaflet and (b) the lower leaflet in the presence of different helices.

As shown in Table I, the order parameter asymmetry induced by the insertion of the helices of Epsin and Sar1p into the membrane is much larger, compared to the insertion of the helix of Arf1. This is likely related to the high efficiency of Epsin and Sar1p in creating membrane deformation. On the basis of above results, we imply that the insertion of amphiphilic helix into the membrane induces an asymmetry in the lipid packing of the membrane’s two leaflets, which further initiate the membrane deformation. Moreover, a relatively shallow insertion depth into the membrane is more efficient in generating high asymmetry in the lipid packing. The extent of the asymmetry in the lipid packing induced by the amphiphilic helix is positively correlated with the efficiency of the helix in producing membrane curvature.

### D. Membrane deformation induced by the insertion of multiple helices

In real experiments, typically multiple proteins cooperate together to create a membrane bud or tube. In order to directly investigate the membrane bending when there are multiple helices inserted into the membrane, the coarse-grained simulation is further performed since it can obtain a much larger space scale and longer time scale.^50–52^ A larger membrane with 4320 lipids is built in the coarse grain simulation to include 20 helices into the systems.

After initially placing multiple helices into the membranes, the membranes are found to be deformed by these helices. Fig. 6 shows the final configurations of multiple amphiphilic helices of Epsin, Sar1p and Arf1 interacting with the membrane. In Fig. 6, the membranes bend toward to the area enriched of the amphiphilic helices, and these amphiphilic helices spread on the area with distinguished membrane deformation. We further characterize the extent of membrane deformation by using the maximum curvature along the membrane that is covered by the amphiphilic helices.^24^ Specifically, the coordinates of the CG beads of the lipid molecules are projected onto the x-z plane, and these projected scattered points are interpolated to estimate the membrane shape. The maximum curvature along the estimated membrane shape is used to characterize the degree of membrane deformation. The time evolutions of the maximum curvature under the insertion of the N-terminal helices of Epsin, Sar1p and Arf1 are plotted in Fig. 7. The averaged value for the estimated curvature radius induced by the membrane insertion of the amphiphilic helices of Epsin, Sar1p and Arf1 are 59 nm, 55 nm, 152 nm respectively (see Table. I), which is in agreement with the inference in the all-atom simulation, namely the N-terminal helices of Epsin or Sar1p induce a much larger membrane curvature than that of the Arf1. Since the experiment conditions contain many more other factors that are not easy to be considered in the molecular dynamics simulation^8–10^, these values are still not able to quantitatively compare with the size of bud or tube induced by Epsin, Sar1p and Arf1 in the experiments. Nevertheless, the general trends are similar between the experiments and the results of all-atom, coarse-grained simulations here, *i.e*., Epsin and Sar1p are more powerful in generating membrane deformations.

**Figure 6.**
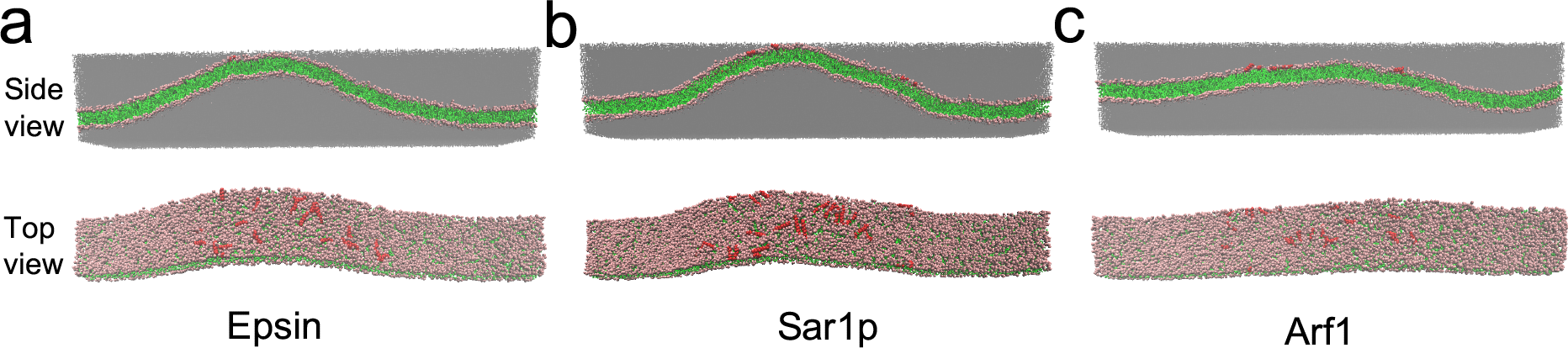
Final configurations of multiple amphiphilic helices of Epsin, Sar1p and Arf1 interacting with the membrane.

**Figure 7.**
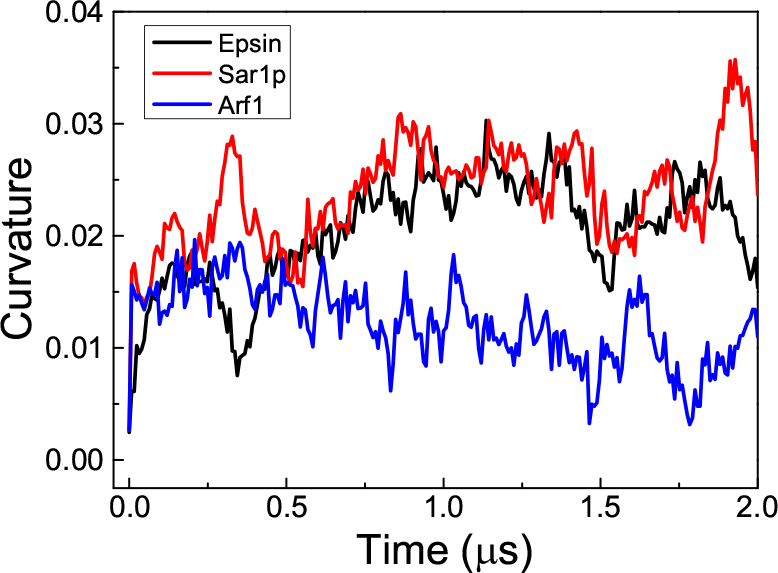
Time evolutions of the mean curvature of the membrane segment enriched of amphiphlic helices of Epsin, Sar1p or Arf1.

## IV CONCLUSION

In summary, we have performed all-atom and coarse-grained molecular dynamics simulations to study the interactions of N-terminal amphiphilic helices of Epsin, Sar1p and Arf1 with the lipid membranes. In all-atom simulations, we find that these amphiphilic helices insert themselves shallowly into the lipid matrix, and induce an asymmetry in the lipid packing of the membrane. Importantly, a shallow rather than a deep insertion of the amphiphilic helix into the membrane is more suitable for generating high asymmetry in the lipid packing. Besides, the induced asymmetry in the lipid packing may be the driven force to deform membranes. Since the helices of Epsin and Sar1p induce a larger asymmetry in the lipid packing than the helix of Arf1, the former two helices could produce a larger membrane deformation than the later one, which is verified in the coarse-grained simulations. Generally, these results are consisted with the experimental results qualitatively that Epsin and Sar1p are more powerful in bending the membrane. In terms of peptide design, a helical peptide with a narrow but clear hydrophobic face seems to be optimized for inducing the membrane deformation. Our findings here can enhance the understanding of the protein-driven membrane remodeling process.

## Acknowledgements

This work is supported by the National Natural Science Foundation of China (No. 91427302, 11474155 and 21604060). We are grateful to the High Performance Computing Center (HPCC) of Nanjing University for doing the numerical calculations in this paper on its IBM Blade cluster system.

